# Dual role of protease activated receptor 4 in acute kidney injury: contributing to renal injury and inflammation, while maintaining the renal filtration barrier upon acute renal ischemia reperfusion injury

**DOI:** 10.1101/540427

**Authors:** Marcel. P. B. Jansen, Nike Claessen, Per W.B. Larsen, Loes M. Butter, Sandrine Florquin, Joris J.T.H. Roelofs

## Abstract

Ischemia reperfusion (I/R) injury triggers the activation of coagulation and inflammation processes involved in the pathophysiology of acute kidney injury (AKI). Coagulation proteases upregulated upon renal I/R injury activate protease activated receptors (PARs), which form an important molecular link between inflammation and coagulation. PAR4 is the major thrombin receptor on mouse platelets, and the only PAR that is expressed on both human and murine platelets. In addition, PAR4 is expressed on other cells including podocytes. We here sought to determine the contribution of PAR4 in the host response to renal I/R injury. Hence, we subjected PAR4 knockout and wild-type mice to renal I/R injury. PAR4 knockout mice exhibited an increased tolerance to renal tubular necrosis and showed a decreased neutrophil influx in response to renal I/R, independent from platelet PAR4. On the other hand, PAR4 deficiency resulted in albumin cast formation in peritubular capillaries and showed a tendency towards albuminuria. Transmission Electron Microscopy revealed an increase in podocyte foot process effacement. Our findings suggest that PAR4 contributes to renal injury likely through facilitating neutrophil migration, independent from platelet PAR4. In addition, PAR4 fulfils an important function in the maintenance of podocyte integrity following renal I/R insult. Subsequently, loss of PAR4 results in albuminuria.

## Introduction

Acute kidney injury (AKI) is a common and costly complication in hospitalized patients, and is independently associated with increased risk of death(1). AKI severity is directly related to patient outcome; even minor changes in serum creatinine predict prognosis in AKI patients after major surgery.(2) Despite the progress in knowledge of the underlying pathophysiology, a recent epidemiological study demonstrated that the incidence of AKI continues to grow.(3) AKI is often triggered by ischemia reperfusion (I/R), which leads to a complex interplay between inflammatory and coagulation processes and renal tissue remodelling(4, 5). More comprehensive understanding of the mechanism underlying these processes is needed to devise strategies to control AKI or accelerate renal recovery. Host-derived coagulation proteases, such as the serine proteases thrombin and factor Xa have been implicated in renal I/R injury(5, 6). Serine proteases regulate haemostasis as well as inflammation and tissue remodelling, thus linking the underlying processes involved in the pathophysiology of AKI(7, 8). Serine proteases elicit cellular effects through cleavage of Protease Activated Receptors (PARs). PARs are a family of seven transmembrane spanning G-protein-coupled receptors that are broadly expressed on immune cells, platelets, and also on renal cells.(9) Upon cleavage, a previously cryptic sequence becomes exposed and acts as a receptor-activated tethered ligand(10), hereby initiating downstream signalling. The PAR family consists of 4 members (PAR1 to PAR4). It has been demonstrated that PAR4 is expressed on podocytes and proximal tubular epithelial cells in both mice and human.(9) Furthermore, PAR4 is the major thrombin receptor on mouse platelets, and the only PAR that is expressed on both human and murine platelets.(11) Although PAR1 and PAR2 have been studied more extensively in context of I/R injury disorders including AKI,(12, 13) the role of PAR4 in the pathophysiology of I/R induced AKI remains largely unknown. Thus, in this study we investigated the effect of PAR4 deficiency upon renal I/R.

## Material and methods

### Animals

Specific pathogen-free C57BL/6J mice were purchased from Harlan Sprague-Dawley (Horst, the Netherlands). PAR4KO mice were purchased as embryos from Mutant Mouse Regional Resource Centers and backcrossed 8 times into the C57BL/6J background in the animal research Institute Amsterdam at the Academic Medical Center. Experimental groups were age-and sex-matched, and housed in the animal research Institute Amsterdam facility under standard care. All experiments were conducted with mice of 10 to 12 weeks of age. The Institutional Animal Care and Use Committee of the Academic Medical Center approved all experiments.

### Murine renal ischemia reperfusion injury

Unilateral renal I/R injury was induced by clamping the left renal artery for 25 minutes followed by a reperfusion phase of 1 day. Renal I/R procedure was performed under general anesthesia (2% isoflurane). For analgesic purposes, mice received a subcutaneous injection of 0.1 mg/kg buprenorphin (Temgesic; Schering-Plough, Brussels, Belgium). The contralateral kidney was used as control. After removal of the clamp, restoration of blood flow was determined by visual inspection. To study the role of PAR4 upon renal I/R injury PAR4 KO (n=8) and WT (n=8) mice were subjected to unilateral renal clamping. To study the role of platelet PAR4 upon renal I/R injury, PAR4KO mice transfused with WT platelets (PAR4KO + WTplt) (n=7) were compared with PAR4KO mice transfused with PARKO platelets (PAR4KO + PAR4KO plt) (n=6). WT mice transfused with WT platelets (WT + WTplt) (n=5) were compared WT mice transfused with PAR4KO platelets (WT + PAR4KO plt) (n=8).

To sacrifice, mice were anesthetized 2% isoflurane followed by cervical dislocation. Kidneys were either snap-frozen in liquid nitrogen or formalin-fixed followed by paraffin embedment.

### Platelet isolation and transfusion

To obtain non-activated platelets for transfusion, blood from PAR4KO mice (n=15) and WT mice (n=15) was collected with a 21-G needle from the inferior vena cava and diluted 4:1 with citrate. Blood was centrifuged at room temperature for 15 minutes at 180 g to obtain platelet rich plasma (PRP). PRP was centrifuged at 250 g for 20 minutes in the presence of acid citrate dextrose, 85 mM Na_3_C_3_H_5_O(COO)_3_, 110 mM Glucose, 65 mM C_6_H_8_O_7_); platelet pellets were washed in 6 mL Buffer A (152 mM NaCl, 13.2 mM NaHCO_3_, 6.16 mM Glucose, 1.1 mM MgCl_2_ 6H_2_O, 2.9 mM KCl, 1mM EDTA, pH6.5). Platelets were re-pelleted and resuspended in 1 mL SSP+ (Storage solution for platelets, Sanquin, Amsterdam, the Netherlands). Platelet count and platelet activation were determined by flow cytometry (FACS Calibur, Becton Dickinson, Franklin Lakes, NJ) using hamster anti-CD61-APC monoclonal antibody (BioLegend, San Diego, CA) and anti-CD62p-FITC (BD Biosciences, San Jose, CA) in accordance with manufacturers’ instructions. Platelet transfusates (200 ul ~3*10^7^/ 20 g BW) were administered directly after isolation via the tail vein, two hours before renal I/R. This number of injected mouse platelets corresponds to approximately 2% of total platelet count(14).

### Enzyme-linked immunosorbent assay

Kidneys were homogenized in a lysis buffer containing 150mM NaCl, 15mM Tris, 1mM MgCl_2_, 1mM CaCl_2_, 1% Triton and 1% protease inhibitors. Concentrations of monocyte chemoattractant protein 1 (MCP-1), keratinocyte-derived chemokine (KC), tumor necrosis factor-α (TNF-α), were measured in kidney homogenates by ELISA according to the instructions of the manufacturer (R&D Systems, Abingdon, UK). Mouse myeloperoxidase (MPO) concentrations were measured by ELISA in kidney homogenates (Duoset DY3667, R&D System, Abingdon). Concentrations of thrombin-antithrombin complexes (TATc) in kidney homogenates were measured according to the manufacturer guidelines using Enzygnost® TAT micro Kit (Siemens Healthcare Diagnostics, Erlangen, Germany). All tissue measurements were corrected for total protein concentration which was measured by incubating 1μL of 10 times diluted homogenates for 30 minutes at 37°C in 500μL of bicinchoninic acid containing 4% of CuSO4, absorbance was measured at 570nm. Urine albumin (Bethyl laboratories) levels were determined by ELISA according to the manufacturer’s instructions.

### (Immuno)histochemistry

Murine renal tissue was fixed and processed as described previously.(15) Paraffin embedded sections were used for periodic acid-Schiff diastase (PAS-D) staining and immunohistochemistry. The degree of tubular damage was assessed on PAS-D-stained 4-µm-thick sections by scoring tubular cell necrosis in 10 non-overlapping high-power fields (magnification 40x) in the corticomedullary junction. The degree of injury was scored by a pathologist in a blinded fashion on a 5-point scale: 0=no damage, 1=10% of the corticomedullary junction injured, 2=10-25%, 3=25-50%, 4=50-75%, 5=more than 75%, as described previously(8, 16).

To determine platelet accumulation in renal tissue, glycoprotein 1bα (GPIbα) antibody, clone SP219, (dilution 1:200, Spring Bioscience) was used and visualized with 3,3-diaminobenzidine. The percentage of positive anti-GP1balpha staining in 5 non-overlapping fields of magnification 20 was quantified using image analysis software Fiji (open-source platform for biological-image analysis). Tissue sections were incubated with specific antibodies for granulocytes (dilution 1:1000 fluorescein isothiocyanate-labeled anti-mouse Ly6G (Lymphocyte antigen 6 complex, locus G) mAb; BD Biosciences–Pharmingen, Breda, The Netherlands) and albumin (1:500, ITK diagnostics), followed by incubation with the appropriate biotinylated secondary antibody and subsequently visualized with 3.3-diaminobenzidine. Ly-6G positive cells were counted in 10 non-overlapping fields (magnification x20) in the corticomedullary junction. The surface percentage of positive albumin staining in 10 non-overlapping fields (magnification x20) was quantified using image analysis software FIJI.

### Ultrastructural analysis

After fixation in Karnovsky buffer (Paraformaldehyde, 8% aq., Glutaraldehyde, 25% aq. 0.2 M Cacodylate, 0.2 M sucrose, distilled H_2_O) for 48 h, the material was post-fixed with 1% osmiumtetroxide, the tissue samples were block-stained with 1% uranyl acetate, dehydrated in dimethoxypropane, and embedded in epoxyresin LX-112. Electron microscopy sections were stained with tannic acid, uranyl acetate, and lead citrate, and then examined using a transmission electron microscope (Philips CM10; FEI). Images were acquired using a digital transmission electron microscopy camera (Morada 10–12; Soft Imaging System) using Research Assistant software (RvC). Podocyte effacement was analyzed in a blinded fashion by an experienced nephropathologist on multiple randomly taken EM images (magnification x4500) in a semiquantitative fashion on a scale of 0-3.

### Quantitative polymerase chain reaction

Total RNA was extracted from snap-frozen renal tissue sections with Trizol reagent (Invitrogen, Life Technologies, Breda, the Netherlands) and converted to cDNA. mRNA level was analyzed by qPCR with SYBR green PCR master mix. SYBR green dye intensity was analyzed with linear regression analysis using LinReg PCR software(17). Specific gene expression was normalized to mouse housekeeping gene cyclophylin G or TATA-box binding protein (TBP). The following murine primer sets were used: cyclophylin G (forward 5’-AAGGGAATGGAAGAGGAGGA-3’ and reverse 5’-CCCTCTGTTGGCCATTGATA-3’); PAR4KO (forward 5’-GATGTTTCCTGGGCTGGG-3’ and reverse 5’-GGTTTTCCCAGTCACGACG-3’); PAR4WT (forward 5’- TGATCCTGGCAGCATGTG-3’ and reverse 5’-TAGGCTCCATTTCTGATCCACC-3’); TBP (forward 5’-GGAGAATCATGGACCAGAACA-3’ and reverse 5’- GATGGGAATTCCAGGAGTCA-3’); MCP-1 (forward 5’-CATCCACGTGTTGGCTCA and reverse 5’-GATCATCTTGCTGGTGAATGAGT-3’); KC (forward 5’-ATAATGGGCTTTTACATTCTTTAACC-3’, and reverse 5’-AGTCCTTTGAACGTCTCTGTCC-3’); TNF-α (forward 5’-CTGTAGCCCACGTCGTAGC-3’, and reverse 5’-TTGAGATCCATGCCGTTG-3’); Neutrophil gelatinase-associated lipocalin (NGAL) (forward 5’-GCCTCAAGGACGACAACATC -3’ and reverse 5’-CTGAACCAATTGGGTCTCGC-3’); Kidney injury molecule-1 (KIM-1) (forward 5’-TGGTTGCCTTCCGTGTCTCT-3’ and reverse 5’-TCAGCTCGGGAATGCACAA-3’)

### Statistical analysis

All data sets were tested for their distribution prior to analyses. Differences between experimental groups were determined using Mann-Whitney U test. Statistical analysis on human data was performed using Kruskal Wallis with Dunn’s post-hoc testing. Correlations were performed using Spearman’s test. All analyses were done using GraphPad Prism version 5.01 (GraphPad Software, San Diego, CA) All data are presented as mean ± SEM (standard error of the mean). A P-value of <0.05 was considered as statistically significant.

## Results

### Protease-activated receptor 4 expression and renal thrombin generation upon renal ischemia/reperfusion injury

To determine renal PAR4 expression under physiological conditions and upon renal I/R injury, we quantified PAR4 mRNA levels in kidney tissue samples from WT and PAR4KO mice. Renal tissue from WT mice express PAR4 mRNA under normal conditions and expression did not change upon renal I/R. PAR4KO mice demonstrated absence of renal PAR4 mRNA expression under normal condition and upon renal I/R injury, demonstrating successful ablation of PAR4 (Fig 1A). To analyze renal thrombin generation, we measured TAT-c in renal tissue homogenates. Here, we show that renal I/R injury results in renal thrombin generation in both WT and PAR4KO mice (Fig 1B). To test the effect of PAR4 deficiency on intra-renal platelet accumulation following renal I/R, we visualized platelets on renal tissue sections. No difference was detected between WT and PAR4KO mice following renal I/R injury (Fig 1C-E).

**Fig 1.**
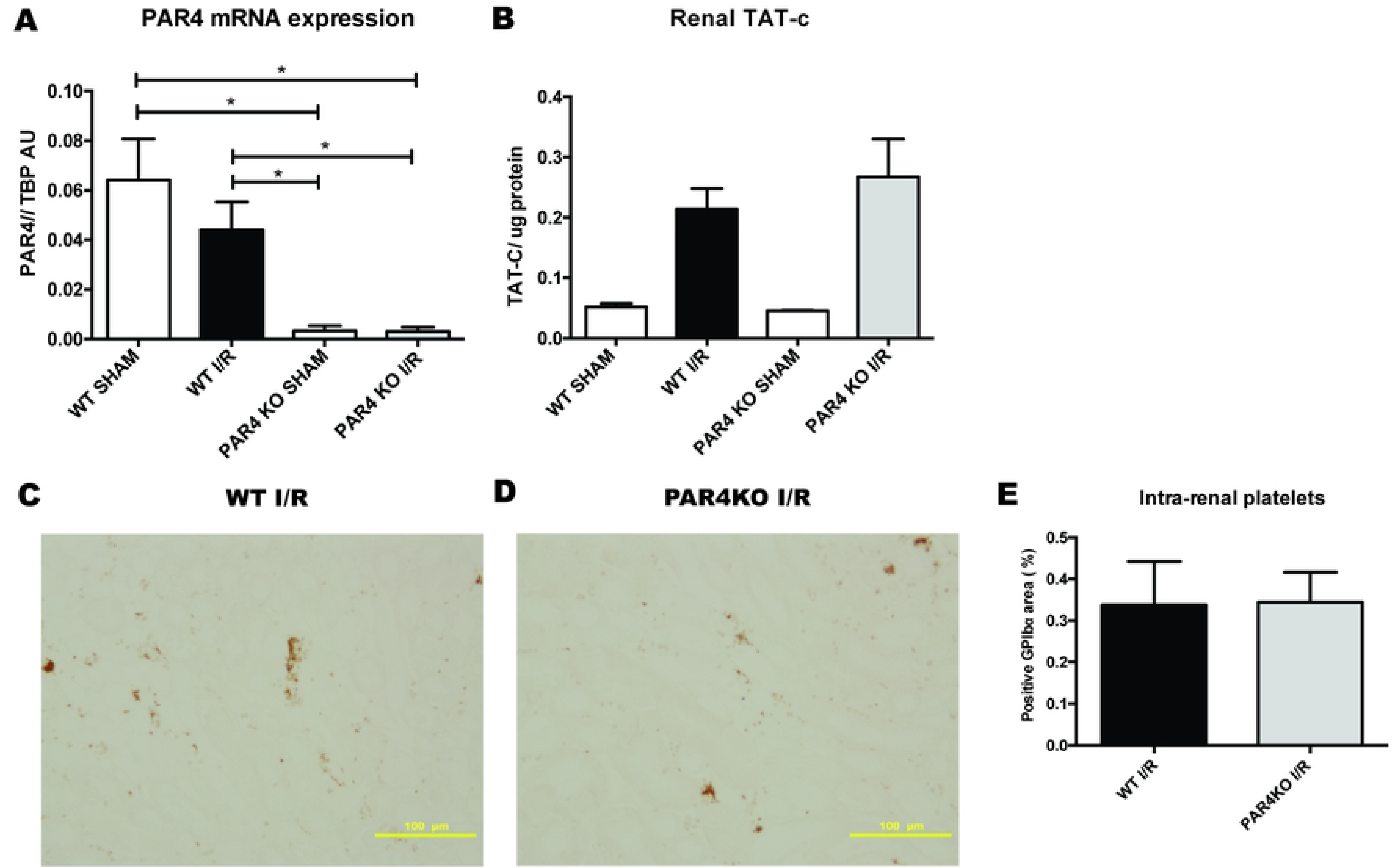
PAR4 expression and renal thrombin generation upon renal ischemia/reperfusion injury. Relative PAR4 mRNA levels were measured in kidneys of WT sham operated mice (white bar, n=8), WT mice 1 day after renal I/R (black bar, n=8), PAR4KO sham operated mice (white bar, n=8) and PAR4KO mice 1 day after renal I/R (grey bar, n=7). Concentrations of thrombin-antithrombin complexes (TATc) were measured in kidneys of WT sham operated mice (white bar, n=2), WT mice 1 day after renal I/R (black bar, n=8), PAR4KO sham operated mice (white bar, n=2) and PAR4KO mice 1 day after renal I/R (grey bar, n=8). Representative pictures of anti-GPIbalpha stained renal tissue sections 20x magnification, scale bar=100µm of WT mice (C) and PAR4KO mice (D), 24 hours following renal I/R. Percentage of positive GP1ba staining in 5 non-overlapping fields (magnification 20x) was quantified using image analysis software FUJI (E) Data are mean±SEM. *: p<0.05

### Genetic ablation of protease-activated receptor 4 decreases renal ischemia/reperfusion injury but increases protein cast formation

To evaluate the functions of PAR4 expression following renal I/R, WT and PAR4KO mice were subjected to renal I/R injury. PAR4 deletion resulted in a modest, but significant decrease of renal injury as assessed by PASD scoring (Figs 2A-C). Of note, PAR4 gene ablation did not result in reduced mRNA expression of kidney injury markers KIM-1 and NGAL (Figs 2D and E). In addition, we found protein casts, present in the tubular lumen of PAR4 KO mice, but absent in WT mice (Figs 2F-J). Taken together, these results demonstrate that PAR4 plays a dual role in the pathophysiology of renal I/R injury: PAR4 is implicated in the processes leading to renal tissue injury, and plays a role in regulating protein filtration and/or reabsorption following renal I/R insult.

**Fig. 2.**
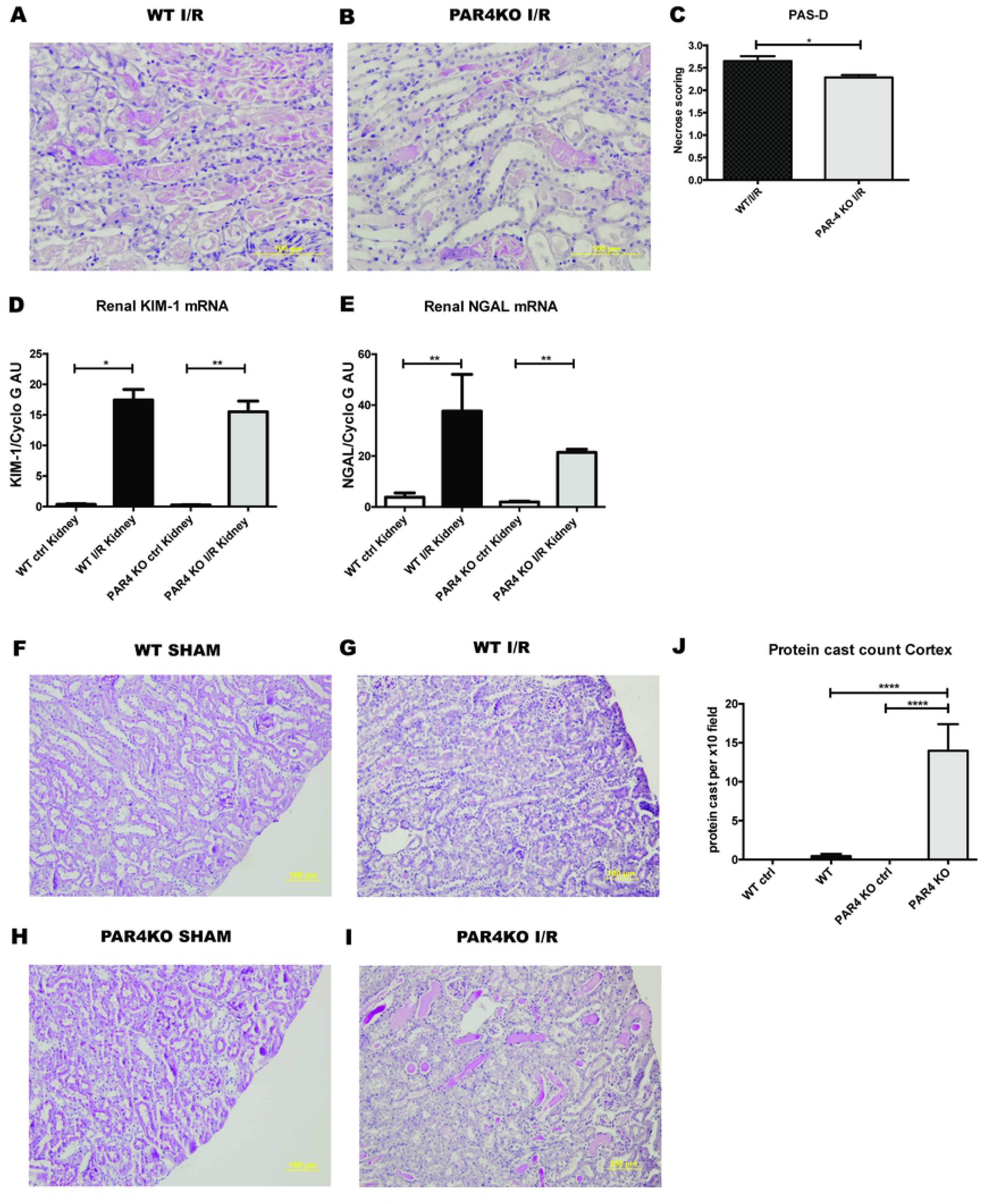
Genetic ablation of PAR4 decreases renal I/R injury but increases protein cast formation. Representative pictures of PAS-D stained renal tissue sections of WT mice subjected to renal I/R with 1 day reperfusion (n=8), 20x magnification, scale bar=100um, (A) and PAR4KO mice subjected to renal I/R with 1 day reperfusion (n=8), 20x magnification, scale bar=100um (B). Score for histopathology of renal tubular damage of WT mice (black bar) and PAR4KO mice (grey bar) subjected to renal I/R injury with 1 day reperfusion (C). Measurement of relative mRNA levels of renal damage marker KIM-1 (D) and NGAL (E) to mouse housekeeping gene cyclophylin G in kidneys of WT sham operated mice (white bar, n=6), WT mice 1 day after renal I/R (black bar, n=8), PAR4KO sham operated mice (white bar, n=8) and PAR4KO mice 1 day after renal I/R (grey bar, n=8). Representative pictures of PAS-D stained renal tissue sections (n=8), 10x magnification, scale bar=100um, of WT mice sham operated (n=8) (F), WT mice after 1 day renal I/R (n=8) (G), PAR4KO mice sham operated (n=8) (H), PAR4KO mice after 1 day renal I/R (n=8) (I). Quantification of positive protein casts in 10 non-overlapping (magnification 10x) was quantified (J). Data are mean ± SEM. *: p<0.05, **: p<0.01,****: p<0.0001

### Protease-activated receptor 4 deletion reduces leukocyte infiltration

Inflammation plays a pivotal role in the pathology of renal I/R injury.(18) Since PAR4 activation modulates several aspects of inflammation including leukocytes recruitment(19), we next explored whether PAR4 deletion decreases inflammation following renal I/R injury. Histological examination of WT and PAR4KO kidneys demonstrated a significant reduction of granulocyte influx in kidneys from PAR4 KO mice (Figs 3A-C). However, enzymatic assay for determining granulocyte activity by measuring MPO did not show differences between WT and PAR4 KO (Fig 3D).

**Fig. 3.**
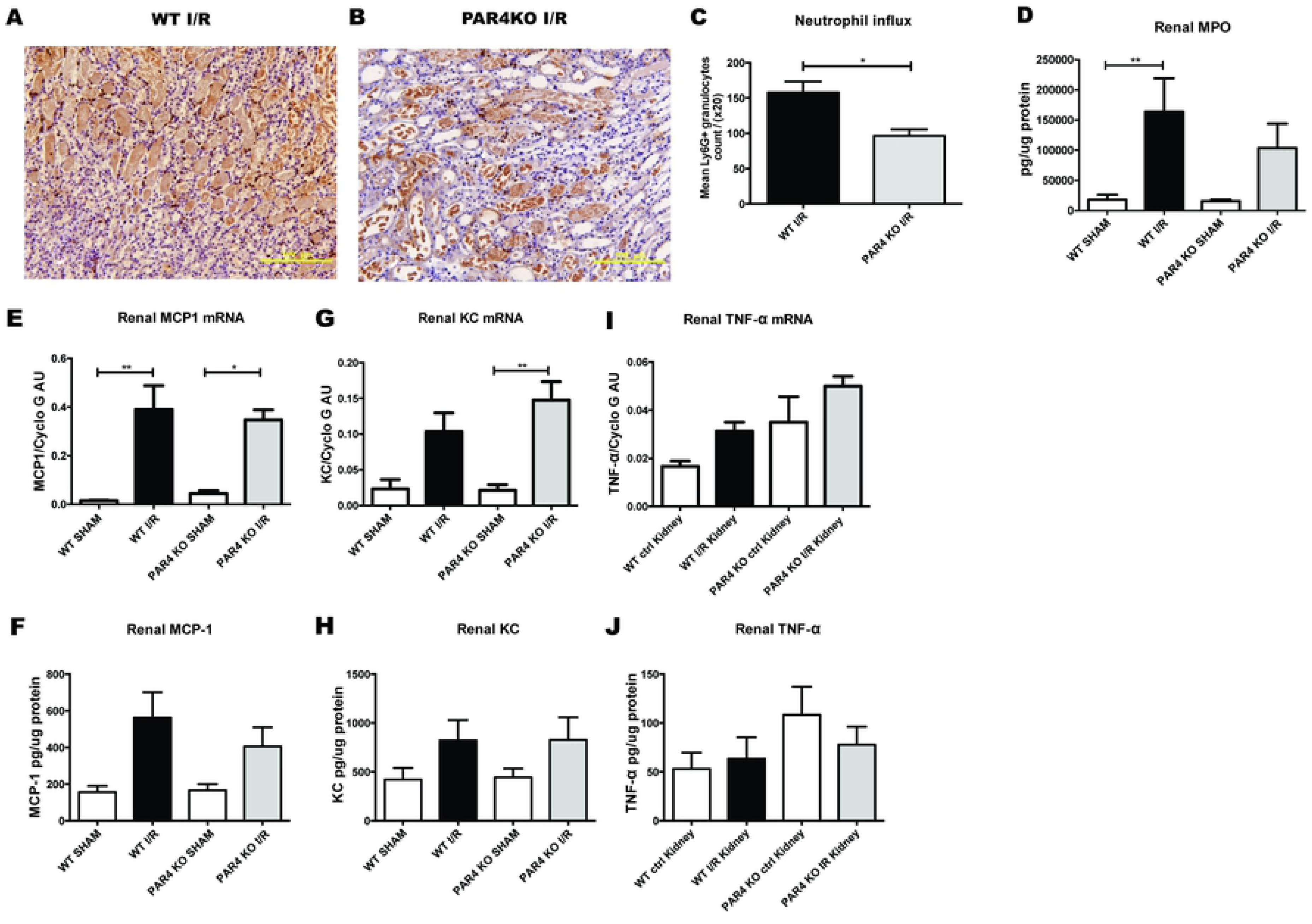
PAR4 deletion reduces leukocyte infiltration. Representative pictures of neutrophil (Ly6-G positive) stained renal tissue sections (20x magnification, scale bar=100µm) of mice WT mice (n=8) (A) and PAR4KO mice (n=8) (B) after 1 day renal I/R. Quantification of positive Ly6G staining in 10 non-overlapping (20x magnification) (C). Concentration of MPO (D) measured by ELISA in kidney homogenates from WT sham operated mice (white bar, n=8), WT mice 1 day after renal I/R (black bar, n=8), PAR4KO sham operated mice (white bar, n=8) and PAR4KO mice 1 day after renal I/R (grey bar, n=8). Measurement of inflammatory cytokine monocyte chemoattractant protein 1 (MCP1) (E & F), keratinocyte chemoattractant (KC) (G & H), tumor necrosis factor-α (TNF-α) (I & J) mRNA expression and protein exposure in renal tissue from WT sham operated mice (white bar, n=8), WT mice 1 day after renal I/R (black bar, n=8), PAR4KO sham operated mice (white bar, n=8) and PAR4KO mice 1 day after renal I/R (grey bar, n=8). Data are mean ± SEM. *: p<0.05, **: p<0.01

To further assess the extent of inflammation, we measured the expression of proinflammatory cytokines MCP-1, KC and TNF-α on mRNA and protein level. PAR4 KO mice showed a trend towards increased TNF-α mRNA expression following renal I/R, while the other cytokines and chemokines were not different (Figs 3 E-J). Since PAR4 is the major thrombin receptor on mouse platelets,(11) we hypothesized that platelet PAR4 might influence renal injury and granulocyte influx in our mouse model. However, transfusion of ~3*10^7^/ 20 g/BW (corresponding to approximately 2% of total platelet count in C57BL/6 mice(14)) PAR4KO platelets and WT platelets in PAR4KO mice or WT mice did not affect renal injury following renal I/R (S1A and S1B Figs). Likewise, transfusion of ~3*10^7^/ 20 g/BW PAR4KO platelets and WT platelets in PAR4KO mice or WT mice did not influence renal granulocyte influx (S1C and S1D Figs). Taken together, these results indicate that murine PAR4 plays an important role in neutrophil tissue influx upon renal I/R injury, which was not influenced by platelet transfusion of approximately 2% PAR4KO or WT platelets of total platelet count.

### Genetic ablation of protease-activated receptor 4 in mice results in increased albuminuria and loss of podocyte structural integrity upon renal ischemia reperfusion

Albuminuria is shown to be an independent mediator of progressive kidney damage.(20) The presence of protein casts in PAR4KO mice upon renal I/R suggests ongoing albumin leakage. To evaluate albuminuria, we collected urine to determine the albumin/creatinine ratio (ACR). In addition, we stained renal sections for albumin to confirm that the observed protein casts consist of albumin. PAR4KO mice show a trend towards increased ACR after I/R (Figure 4A). Under normal conditions, WT and PAR4KO mice did not show any albuminuria. Albumin staining on renal sections demonstrates the presence of albumin in the protein casts (Figure 4B, C), which is significantly increased in PAR4KO compared to WT mice following renal I/R injury (Figure 4D).

**Fig. 4.**
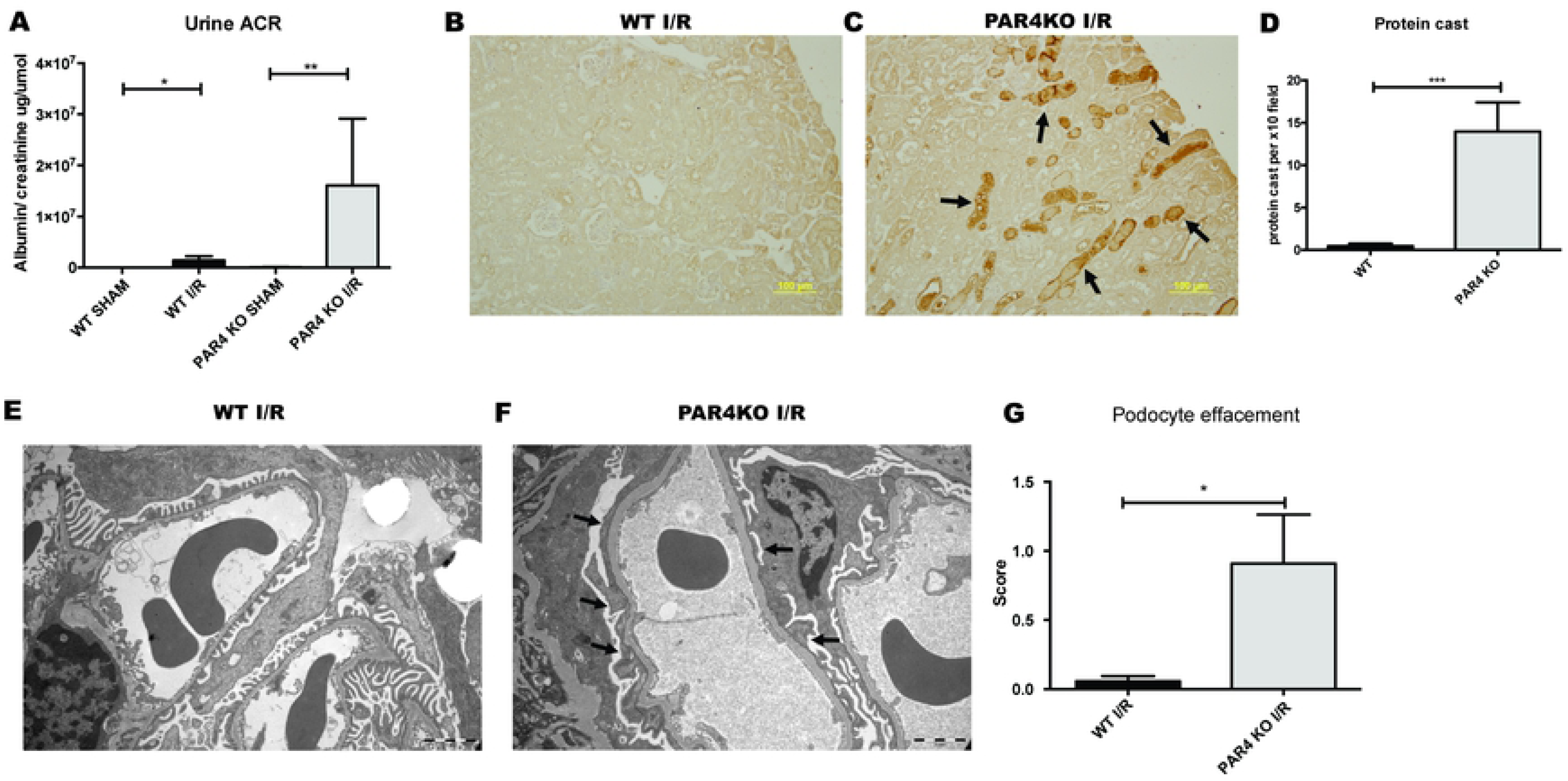
Genetic ablation of PAR4 in mice results in increased albuminuria and loss of podocyte structural integrity upon renal I/R. Measurement of albumin creatinine ration in WT sham operated mice (white bar, n=7), WT mice 1 day after renal I/R (black bar, n=8), PAR4KO sham operated mice (white bar, n=8) and PAR4KO mice 1 day after renal I/R (grey bar, n=8) (A). Representative pictures of albumin staining in WT (B) and PAR4KO (C) mice 1 day after renal I/R. Quantification of positive albumin casts in 10 non-overlapping (magnification 10x, scale bar= 100µm) (D). Representative electron micrographs of podocytes surrounding a glomerular capillary in WT mouse (E) and PAR4KO (F) mouse (magnification 4500x, scale bar= 2µm) after 1 day renal I/R (arrows point to podocyte effacements). Podocyte effacement semi-quantitative score on a scale of 0-3, on multiple randomly taken EM images (magnification 4500x) in WT and PAR4KO mice after 1 day renal I/R (G). Data are mean ± SEM. *: p<0.05, **: p<0.01, ***: p<0.001

To investigate whether the origin of albuminuria lies in the glomerular filter, we assessed the glomeruli of PAR4KO and WT mice by transmission electron microscopy to investigate ultrastructural differences. No structural podocyte abnormalities were encountered in both sham-treated WT and PAR4 KO mice (S2 Fig). However, upon renal I/R PAR4KO deficient mice demonstrated a significantly increased loss of podocyte structure integrity as reflected by extensive foot process effacement (Figure 4E-G). Since the origin of albumin leakage can also be found in the proximal tubular reabsorption machinery, in which endocytic receptor megalin plays an important role in albumin reabsorption,(21, 22) we evaluated the tubular expression of albumin reabsorption receptor megalin by immunostainings. No differences were found (data not shown). Taken together these results imply that PAR4 is implicated in maintaining podocyte integrity upon renal I/R insult.

## Discussion

AKI is a common and costly complication frequently caused by ischemia reperfusion (I/R) injury,(1) leading to a complex interplay between inflammatory and coagulation processes and renal tissue remodelling.(4, 23) Serine proteases such as thrombin and factor Xa regulate haemostasis as well as inflammation and tissue remodelling, and have been associated with renal I/R injury(5, 6). The cellular effects of serine proteases are elicited by cleaving of PAR’s, comprising of PAR1-4(10) which are widely expressed on myeloid cells and various renal cells.(9) Recently it has been shown that PAR4 deficiency offers protection against acute I/R injury in cerebral and heart tissue.(24, 25) To date the role of PAR4 upon renal I/R is unknown. Here we sought to determine the role of PAR4 in the host response to renal I/R injury. Our main finding is that genetic ablation of PAR4 gene decreases renal necrosis and leukocyte influx in the corticomedullary region, independent from platelet PAR4. In addition, following renal I/R injury, PAR4 deficiency resulted in intra-tubular albumin casts and albuminuria, likely through loss of podocyte structural integrity. Taken together these results suggest that PAR4 is involved in the pathophysiology of renal I/R injury enabling leukocytes influx into renal tissue. On the other hand, PAR4 also fulfils a functional role in the maintenance of podocyte structure upon renal I/R injury insult.

In a study of Kolpakov et al, it was shown that PAR4 is involved in pathophysiology of myocardial I/R.(24) They showed that PAR4 expression in WT mice is upregulated in heart tissue following acute myocardial I/R and demonstrated that ablation of the PAR4 gene results in smaller heart infarct size and decreased myocyte death. In correspondence with this model, the study of Mao et al demonstrated that in a mouse model of cerebral I/R injury, PAR4 deficiency attenuated I/R injury.(25) In contrast to the study of Kolpakov et al, WT mice in our renal I/R mouse model did not show increased renal PAR4 expression upon I/R. In line with Kolpakov et al. and Mao et al., we demonstrated that PAR4KO mice show less renal tissue injury following renal I/R when compared to WT mice. However, albeit significant, the difference was modest between PAR4KO and WT mice, which may explain why mRNA expression levels of kidney damage markers KIM-1 and NGAL did not differ between WT mice and PAR4KO mice.

PAR4 has been shown to be involved in acute inflammatory responses.(26) Vergenolle et al. showed that PAR4 agonists cause neutrophil rolling and adherence, indicating that PAR4 activation contribute to the recruitment of neutrophils.(19) Migration of neutrophils into renal parenchyma is a significant component in the pathophysiology of renal I/R injury.(18, 27) Neutrophils can aggravate kidney injury by releasing proteases such as myeloperoxidase, reactive oxygen species or NETS(16, 28, 29). In this study, we show that PAR4 deficient mice demonstrate less neutrophil accumulation in the renal parenchyma, indicating that PAR4 plays a role in neutrophil recruitment following renal I/R. To note, PAR4KO did not impact the expression of pro-inflammatory cytokine MCP-1 or KC upon renal I/R, suggesting that PAR4 is more involved in the processes facilitating neutrophil rolling and adherence rather than the renal inflammatory cytokine response. Activated platelets play an important role in facilitating granulocyte influx upon AKI(30, 31) by forming a bridge to the endothelial wall, stimulating granulocyte rolling and adherence.(32) Murine platelets are dependent on PAR4 for thrombin-induced signaling and subsequent activation.(33) We therefore speculated that platelet PAR4 is important for thrombin dependent platelet activation upon AKI and subsequently neutrophil tissue influx and renal injury in our model. However, PAR4 deficiency did not influence platelet accumulation upon renal I/R. In accordance with the study performed by Huo et al. showing that injection of ~3*10^7^/ 20 g/BW activated WT platelets increased leukocyte arrest on the surface of atherosclerostic lesions, we transfused PAR4KO and WT mice with ~3*10^7^/ 20 g/BW PAR4KO or WT platelets prior to renal I/R injury. However, ~3*10^7^/ 20 g/BW corresponding to approximately 2% of total platelet count did not affect renal injury or granulocyte renal influx. This suggests that murine platelet PAR4 is not implicated in the pathophysiology of AKI in our model. However, it may also indicate that more than 2% transfused PAR4KO or WT platelets of total platelet count in mice is needed to affect the neutrophil influx and renal tissue injury. Taken together these results suggest that PAR4 is involved in the pathophysiology of renal injury, possibly through facilitating neutrophil tissue influx, independent of renal cytokine release.

Madhusudhan et al. demonstrated that PAR4 is expressed on podocytes in both mice and human(9, 34), however its function is unknown so far. Podocytes are essential in the maintenance of an intact glomerular filtration barrier and podocyte stress and/or injury is a major cause of albuminuria(35, 36). Strikingly, we here demonstrated for the first time that PAR4 deficiency results in albumin cast formation in the peritubular capillaries. In addition, PAR4 deficient mice show a tendency towards elevated albumin levels in urine following renal I/R, hinting at a problem with the glomerular filtration barrier. Under normal conditions, podocyte structure is characterized by their interdigitated foot processes that are wrapped around the glomerular capillaries and form filtration slits which are bridged by the slit diaphragm.(37) Upon stress or injury the podocytes’ structure can be altered, leading to podocyte foot process effacement, which causes albuminuria.(38) In this study, transmission electron microscopy of the glomeruli revealed increased levels of podocytes foot process effacement in PAR4 deficient mice subjected to renal I/R, suggesting that PAR4 is involved in maintenance of a normal structure of podocytes upon I/R injury. Loss of podocyte PAR4 makes podocytes more susceptible to structural changes upon I/R insult, resulting in increased albumin leakage.

In conclusion, our data shows that PAR4, is implicated in the pathophysiology of renal injury, likely through facilitating neutrophil influx. In addition, PAR4 is involved in maintaining podocyte integrity following renal I/R insult. To what extent platelet specific PAR4 is implicated needs to be further explored in future studies.

## Supporting information

**S1 Fig.** Representative electron micrographs of podocytes surrounding a glomerular capillary in WT mouse (E) and PAR4KO (F) mouse (magnification 4500x, scale bar= 2µm) sham-operated (arrows point to podocytes).

**S2 Fig.** Representative pictures of PAS-D stained renal tissue sections (magnification 40x, scale bar= 50 um) and score of histopathology of renal tubular damage of: PAR4KO mice transfused with ~3*10^7^/ 20 g/BW WT platelets (black bar) or PAR4KO platelets (grey bar) (A), WT mice transfused with ~3*10^7^/ 20 g/BW WT platelets (black bar) or PAR4KO platelets (grey bar) (B). Representative pictures of neutrophil (Ly6-G positive) stained renal tissue sections (magnification 40x, scale bar=50 µm) and quantification of positive Ly6G staining in 10 non-overlapping high power fields (magnification 40X) in PAR4KO mice transfused with ~3*10^7^/ 20 g/BW WT platelets (black bar) or PAR4KO platelets (grey bar) (C), WT mice transfused with ~3*10^7^/ 20 g/BW WT platelets (black bar) or PAR4KO platelets (grey bar) (D).

